# The fate of pollen in two morphologically contrasting buzz-pollinated *Solanum* flowers

**DOI:** 10.1101/2024.09.01.610688

**Authors:** Christian A. Vasquez-Castro, Elodie Morel, Bernardo Garcia-Simpson, Mario Vallejo-Marín

**Affiliations:** Department of Ecology and Genetics, Uppsala University, Uppsala, 752 36, Sweden; Department of Biology, Faculty of Science, Universidad Nacional Agraria La Molina, Lima, 150 24, Peru; Institute of Biology, University of Neuchâtel, 2000, Neuchâtel, Switzerland; Círculo de Investigación en Taxonomía, Florística y Ecología Vegetal, Department of Biology, Faculty of Science, Universidad Nacional Agraria La Molina, Lima, 150 24, Peru

**Author notes:** Co-first authors (Christian A. Vasquez-Castro, Elodie Morel, Bernardo Garcia-Simpson).

**Keywords:** Buzz pollination, floral morphology, heteranthery, pollen carryover, pollen transfer and deposition, pollen fates

## Abstract

Pollen transfer efficiency (PTE) and pollen deposition patterns on a pollinator’s body significantly influence plant reproductive success. However, studies on pollen fates (i.e., the destination of pollen grains after anther dehiscence) in animal-pollinated species offering pollen as the sole reward are limited. This study investigated pollen fates in two nectarless, buzz-pollinated *Solanum* species with contrasting floral morphology. Experimental trials were conducted involving one pollen donor and four recipient flowers of *Solanum rostratum* Dunal or *S. dulcamara* L., using captive *Bombus terrestris* L. as pollinator. The number of pollen grains remaining in the anthers, deposited on stigmas, placed on the pollinator, and falling to the ground was quantified. Both species produced a relatively high number of pollen grains as expected for buzz-pollinated plants. Pollen deposition curves followed exponential decay patterns, with a higher number of pollen grains deposited for *S. dulcamara*, and a similar rate of decline for both species. PTE was similar between species (0.86 % vs. 1.00 %, for *S. rostratum* and *S. dulcamara,* respectively) but these values could be 25 % higher if we were to measure pollen deposition in 20 recipients rather than four as in the present study. Although both species had similar PTE values, their pollen fates differed: pollen was mainly lost on the ground in *S. rostratum* and due to bee grooming in *S. dulcamara*, potentially explained by their different floral architectures. These findings suggest that species with different flower morphology could exhibit different pollen fates without impacting pollen transfer to conspecific stigmas.

## Introduction

Pollen transfer from anthers to stigmas is a major determinant of plant fitness (Minnaar et al. 2019; Christopher et al. 2020). Plants produce a large number of pollen grains per flower (Cruden 2000), yet very few pollen grains produced in anthers ever make it to their destination. For instance, it is estimated that, on average, only 2.3 % of pollen grains removed from animal-pollinated, nectar-rewarding plants with monads (i.e., pollen grains released as single units) reach the stigma of conspecific individuals (Harder & Thomson 1989; Johnson & Harder 2023; Supplementary material from Johnson & Harder 2023).

Pollen fates can be decomposed in three phases: pollen production and presentation, pollen transfer, and pollen germination (Minnaar et al. 2019). Among these, pollen transfer, the pathway between pollen removal and pollen deposition, is highly stochastic and mediated by the behaviour of pollinators in animal-pollinated plants (Thomson & Plowright 1980; Richards et al. 2009) (Fig. 1). Despite the growing interest in its study since Thomson and Plowright (1980) and its importance for plant reproductive success, conducting studies on pollen transfer remains challenging, and much past research has focused on species with pollen aggregated in pollinia, such as orchids. This is largely due to the technical difficulties involved in tracking individual pollen grains across the different stages of the pollination process. The pathway followed by pollen is especially hazardous for plants giving pollen as the only reward, as floral visitors actively collect pollen grains during floral visitation and might further reduce the pool of pollen remaining for fertilizing other flowers (Luo et al. 2008; Vallejo-Marín et al. 2009).

**Figure. 1.**
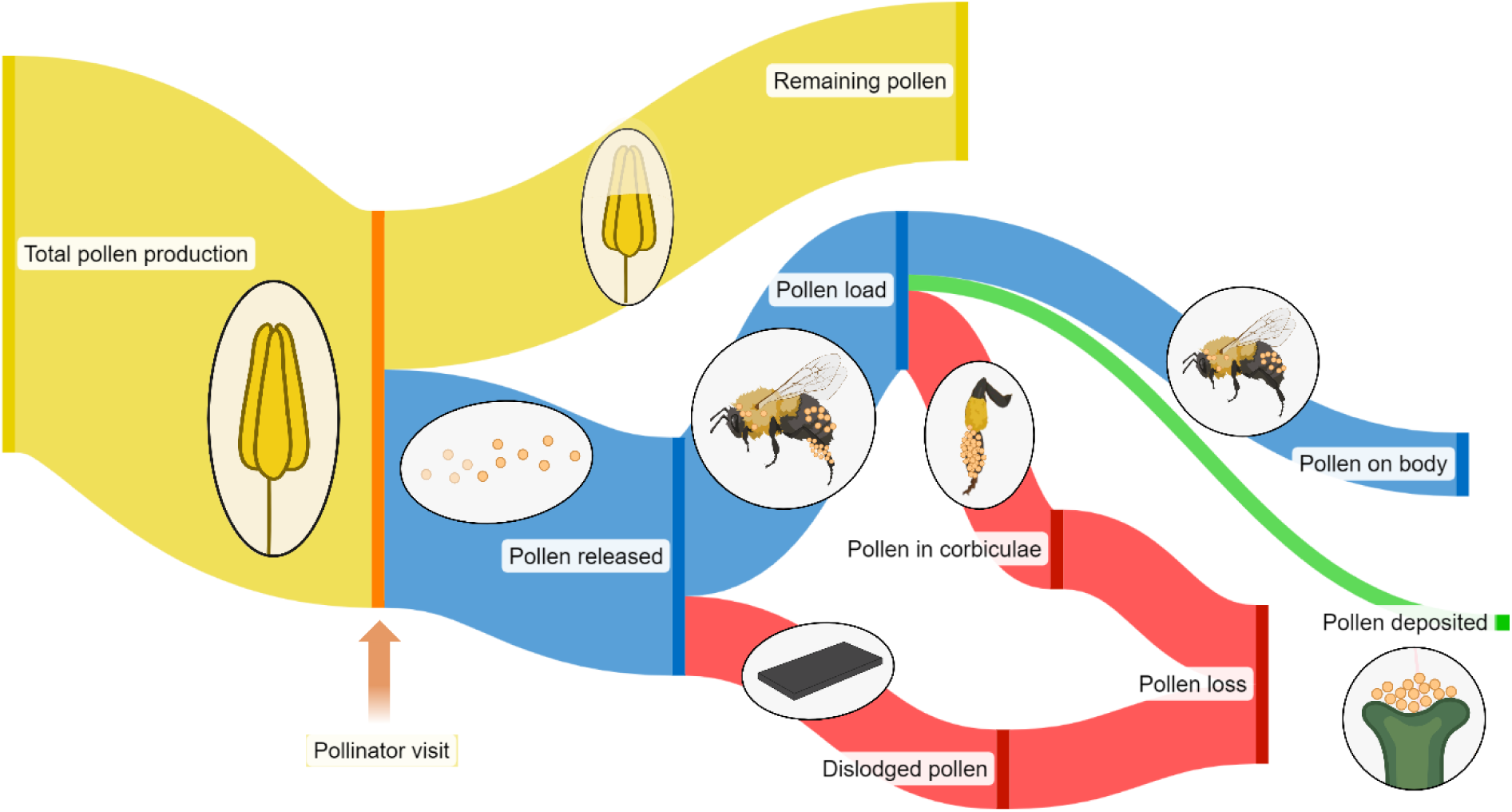
Pathways depicting the fates of pollen in a bee pollinated species such as the ones studied here. The diagram shows pollen contained inside the anthers (yellow); pollen removed that can be potentially transferred to conspecific stigmas (blue); pollen lost from the fertilization process, including dislodged pollen that falls to the ground and pollen stored by the bee (e.g., in corbiculae) and thus unable to contribute to fertilization (red); and pollen deposited on stigmas (green). Created with BioRender.com.

Pollen transfer efficiency (PTE) measures the proportion of pollen removed from the anthers that successfully reaches conspecific stigmas and is consequently available for potential ovule fertilization (Johnson & Harder 2023, Johnson et al. 2005). In animal-pollinated species, PTE usually decreases as the amount of pollen removed per visit increases (Harder & Thomson 1989; Harder 1990, Thomson et al. 2000). This gives a fitness advantage to plants with gradual pollen release or pollen packaging strategies, i.e., mechanisms that divide pollen production in different “packages” (inflorescences, flowers, etc.) which do not necessarily release pollen simultaneously (Harder & Thomson 1989). The Pollen Presentation Theory (Thomson 2006) suggests that natural selection via male fitness should favour traits that regulate pollen dispensing (i.e., restriction of the amount of pollen released by one flower in a single pollinator visit) (Harder & Thomson 1989; Thomson 2006).

In addition to PTE, a key characteristic affecting the fate of pollen and its consequences for plant fitness is how the number of pollen grains deposited on stigmas decreases in a consecutive sequence of recipient flower (i.e., the shape of its pollen deposition curve, referred to as pollen carryover curve; Thomson & Plowright 1980). Pollen deposition curves tend to show an exponential decay pattern, but several studies have found longer tails than expected under an exponential model (i.e., leptokurtic distribution) due to long dispersal events (Lertzman 1981; Price & Waser 1982; Morris et al. 1994; Santa-Martinez et al. 2021). The steepness of the curve varies with pollinator type (Castellanos et al. 2003; Santa-Martinez et al. 2021), type of flower, and reward availability in the recipient flower (Thomson & Plowright 1980). Deposition curves have been studied in hummingbird pollinated *Ipomopsis* and *Castilleja* (e.g., Lertzman 1981; Price & Waser 1982; Morris et al. 1994), and in bee-pollinated species (e.g., Thomson & Plowright 1980; Thomson 1986; Rademaker et al. 1997; Castellanos et al. 2003; Santa-Martinez et al. 2021). These studies have shown that grooming behaviour plays a crucial role in determining the steepness of the curve for bee-pollinated species. However, pollen deposition curves have rarely been estimated for nectarless plants that use pollen as the only reward to floral visitors. Given that in these plants pollen serves both as a vehicle for male gametes and simultaneously as a reward to potential pollinators (Vallejo-Marín et al. 2009), pollen deposition curves could show a particularly steep decline if pollinators are efficient at grooming pollen following a visit.

The present work characterizes and compares the production, release, loss and transfer efficiency of pollen (pollen fates) of two nectarless, bee-pollinated *Solanum* (Solanaceae) species *(S. dulcamara* and *S. rostratum*). As most of the other ∼1,200 described *Solanum* spp. (Hilgenhof et al. 2023), the species studied here are buzz-pollinated, i.e., they are pollinated by pollen-foraging bees that use vibrations to harvest pollen (Buchmann 1983; Vallejo-Marín & Russell 2024). Some buzz-pollinated species release pollen gradually, acting as a pollen dispensing mechanism that could increase PTE (Harder & Barclay 1994; Harder & Thomson 1989, Thomson 2006, Kemp & Vallejo-Marín 2021). Both *S. rostratum* and *S. dulcamara* have bisexual, nectarless flowers, with five poricidal stamens (i.e., tube-like anthers that open through small apical pores at the tip). They differ markedly in their stamen shape and architecture (i.e., the shape and arrangement of anthers within the flower). *S. rostratum* flowers have two morphologically and functionally distinct sets of stamens in the same flower (i.e., heteranthery): one set is composed of four smaller, bright yellow ‘feeding’ anthers centrally located, and the other set includes a single larger, yellow to brown ‘pollinating’ anther that is laterally displaced, opposite to the style (Fig. 2B) (Vallejo-Marín et al. 2009). This stamen dimorphism results in pollen from the pollinating anther being placed in harder-to-groom ‘safe sites’ of the pollinator’s body, reducing pollen loss by grooming, and increasing their chance to reach the stigma (Vallejo-Marín et al. 2009). Individuals of *S. rostratum* are enantiostylous, i.e., each plant produces two types of flowers with the style deflected either to the right- or left-hand side, which reduces anther-stigma interference and promotes pollen transfer among flower morphs (Jesson & Barrett 2002). In contrast, *S. dulcamara* flowers have monomorphic anthers that are fused into an anther cone which surrounds the centrally-placed pistil (Fig. 2A) (Glover et al. 2004). These morphological differences might affect pollen fates through their influence on pollinator grooming and capacity to remove pollen from the anthers. Studying two morphologically distinct species enables the quantification of pollen fates and deposition curves, while also allowing for first explorations of how flower architecture could influence pollen fates and bees’ capacity to remove pollen from the anthers of buzz-pollinated plants.

**Figure 2.**
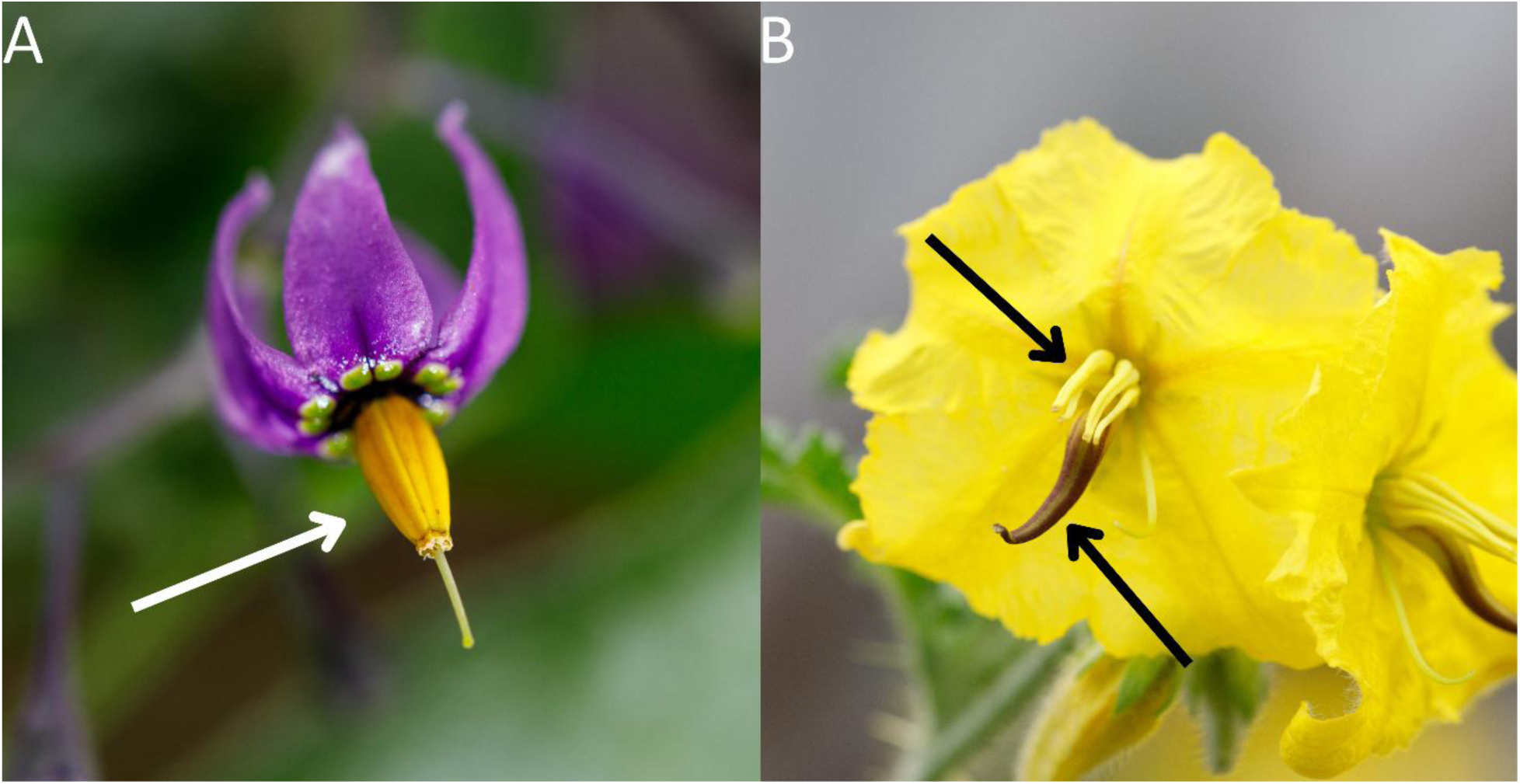
(A) Flower of *S. dulcamara*, arrow indicates fused anthers (anther cone). (B) Flower of *S. rostratum*, arrows indicate the feeding (above) and pollinating anthers (below). Photo credits: M. Vallejo-Marín.

To investigate pollen fates in these two species, a sequence of experimental flowers was exposed to visits by a generalist buzz-pollinating bee, *Bombus terrestris* (Apidae). The specific objectives were: (1) To quantify pollen fates and PTE of two *Solanum* species with contrasting size and floral architecture (*Solanum rostratum* Dunal and *S. dulcamara* L.), and (2) to estimate the pollen deposition curves for both species. We predicted that both species would show higher PTE than the 1-2 % typically reported for nectar-rewarding plants with monads, due to their gradual pollen release strategy. Additionally, pollen deposition curves for both species were expected to follow an exponential decay pattern across recipient flowers. Despite these similarities, distinct pollen fates and steepness of the deposition curves were expected between the species due to variations in floral morphology and pollen placement strategies. Specifically, it was expected that *S. rostratum* would display a less steep pollen deposition curve and lower pollen loss coupled with higher PTE, due to stamen dimorphism. In contrast, it was predicted that *S. dulcamara* would exhibit a steeper pollen deposition curve, higher pollen loss, and lower PTE, likely due to its lack of a specialized stamen configuration making pollen more susceptible to removal during pollinator grooming.

## Materials and methods

### Plants

*Solanum dulcamara* is a perennial vine native to Eurasia and introduced into North America (Knapp et al. 2013). Its flowers are actinomorphic and relatively small (1.5-2 cm diameter). In Europe, *S. dulcamara* is visited by different species of bumblebees (Free 1970) and is self-compatible, but its mating system is unknown. Conversely, *S. rostratum* is an annual herb native to Mexico and perhaps southern USA, and introduced in disturbed habitats around the world (Zhao et al. 2013). Flowers are medium-sized (2.3-3.5 cm diameter). In its native and introduced range, *S. rostratum* is visited by different types of bees, including bumblebees (*Bombus* spp.) (Bowers 1975; Solís-Montero et al. 2015). Genetic estimation of outcrossing rates indicates that *S. rostratum* is predominantly outcrossing (*t* = 0.70) in both native and introduced ranges (Vallejo-Marín et al. 2013; Zhang et al. 2017). For our study, plants of both species were grown from field-collected seeds in glasshouses at Uppsala University. For *S. rostratum,* seeds were used from accessions 10s71-5, 10s74, 10s77, 10s79, and 10s80 (San Miguel de Allende, Guanajuato, Mexico), and 11s182, 11s188, 11s191, 11s205, and 11s207 (Dolores, Guanajuato, Mexico). For *S. dulcamara,* seeds were used from accessions 22-UPP-1 and 22-UPP-2 collected at Uppsala. Seeds were germinated as described in Vallejo-Marín et al. (2014), grown at 16h/20°C light and 8h/16°C dark cycles, and fertilised with general purpose liquid fertiliser.

### Pollinators

To evaluate pollen transfer, we employed *Bombus terrestris*, a generalist, buzz-pollinating bumblebee that is widely distributed in Europe and commonly used to supplement pollination of soft-fruit crops. *Bombus terrestris* visits *S. dulcamara* in the native range and is similar in size and behaviour to other bumblebees visiting *S. rostratum* in the Americas (Bowers 1975; M. Vallejo-Marín, Uppsala University, Sweden, pers. obs). Workers from two commercial colonies (Koppert Biological Systems, Helsingborg, Sweden; and Biobest Group, Westerlo, Belgium) were used. Colonies were maintained on 2 M solution of sucrose available from artificial feeders within enclosed foraging arenas (82×60×60 cm), follow (Russell et al. 2017). Twice a week, bees were also provided with approximately 10 g of “pollen dough” made of pulverised honeybee-collected pollen (Koppert Biological Systems, Helsingborg, Sweden) mixed with sucrose solution and placed inside the colony. The foraging arenas were kept at room temperature and illuminated with an LED ceiling panel set to a 14 h:10 h light:dark cycle. Before the experiment, bumblebees were conditioned to visit *Solanum* flowers by allowing them to forage freely on flowers placed in the foraging arena. This conditioning phase was never done on the same day of the experimental trials to avoid bees carrying over pollen grain collected during the conditioning phase to the experimental trials.

### Experimental trials

Before each trial, fresh flowers were collected from the greenhouse each morning before experiments and carefully transported in floral foam (OASIS Floral Products, Washington, UK) within a closed plastic box. Each trial consisted of one donor flower from which pollen could be removed by the bee, and four recipients (R1-R4) that were experimentally prevented from releasing pollen by sealing the anther pores using a small drop of PVA glue (Elmer’s Products, Westerville, USA) under a stereoscopic microscope. A similar amount of glue was applied just below the anther pore of the donor flowers as a sham treatment. After the glue dried, the pedicel of each flower was mounted on a bamboo stick with Blu-Tack adhesive gum (Bostik, Staffordshire, UK) and placed in a plastic tube containing floral foam. Every flower was placed vertically, with the anthers facing forward at a 90° angle to the ground. This placement was chosen to ensure a consistent positioning throughout the trials, despite not reflecting the natural orientation of flowers in an inflorescence. A black plate was placed under the donor flower to recover pollen falling from the flower (dislodged pollen). The pollen lost on the ground or in the air after a bee departed from the donor flower (transport loss) was not quantified. To control for differences in pollen transfer within and between morphs of enantiostylous *S. rostratum*, all recipients were set to have the reciprocal mirror-image morph of the donor flower (i.e., the left-hand side morph for all recipients if the donor was a right-hand flower, and vice versa).

For each trial, one donor flower was placed in the centre of the foraging arena and a single bee was allowed to enter the arena. Immediately after the bee landed on the donor flower, the first recipient flower (R1) was placed two to five centimetres next to it and the donor flower was removed from the arena as soon as the bee departed from it. The procedure was followed similarly with the consecutive recipients (R2, R3 and R4). If after 10 minutes the bee had not landed on the donor flower, the trial was ended and the bee removed. A total of nine and six trials were successfully completed up to the fourth recipient for *S. dulcamara* and *S. rostratum*, respectively, resulting in 60 recipient flowers evaluated across 15 trials.

All trials were video recorded using a camera (Canon EOS M50, Tokyo, Japan) placed right outside of the arena. Recordings were used to calculate the duration of each flower visit. Temperature, humidity of the arena and time of the day were also recorded.

### Pollen sampling and counting

Immediately after each trial, the bee was captured in a plastic tube and put in the freezer for five to eight minutes to anesthetize it. Once immobilized, the bee was placed under a stereoscopic microscope to remove pollen from its corbiculae. The pollen was placed on a fuchsin jelly cube (32 mm^3^) (Kearns & Inouye 1993). Pollen remaining on the body was collected with a separate jelly cube. Each sample was stored in individual 1.5 mL Eppendorf tubes. The stigmas of the four recipients were individually cut and placed on jelly cubes. Dislodged pollen was collected from the black plate using 1-2 cubes of jelly, and anthers from the donor flower were cut and preserved in 1ml of 70 % ethanol. Each fuchsin jelly cube containing a stigma sample was mounted on a slide and examined under a light microscope at 400x magnification for pollen grain counting. To process the remaining pollen samples, 500 μL of 70 % ethanol was added to tubes containing pollen from the ground, corbiculae, and body. Donor anthers were subjected to sonication using an Elmasonic S40H ultrasonic cleaner (Elma Schmidbauer GmbH, Singen, Germany), followed by staining pollen grains with 200 μL of lactoglycerol-aniline blue (Kearns & Inouye 1993). For pollen grain counting, an aliquot was taken from each tube and placed onto a haemocytometer chamber until 200 pollen grains were counted. To estimate the number of pollen grains, the combined volumes of ethanol, jelly, and aniline used in their respective cases were estimated.

For each trial, the following variables were counted: (C) number of pollen grains remaining in the donor anthers, (D) number of pollen grains that fell on the ground (dislodged from the pollinator), (F) number of pollen grains on the bee’s corbiculae and (G) on the bee’s body, and (H_1-4_) the number of pollen grains deposited on the stigma of each recipient. From these variables, the following derived variables were also calculated: total pollen production (A = C + D + F + G + H), pollen removed (B = F + G + H + D), pollen collected (E = F + G + H), pollen lost from the pollination process (I = D + F), and the sum of the number of pollen grains deposited on the four recipient flowers for each trial (J = H_1_ + H_2_ + H_3_ + H_4_). Each of the variables C, D, F, G, and J was multiplied by 100/A to express values as percentages of total pollen production. Additionally, variables D, F, G, and J (excluding C) were multiplied by 100/B to represent percentages of pollen removed from the donor. In particular, J × 100/B is defined as the Pollen Transfer Efficiency (PTE), calculated for each trial.

### Statistical analyses

To determine whether the proportions of pollen grains deposited on different destinations (body, G; corbiculae, F; ground, D; or recipient stigmas, J) differed between species, generalized linear models (GLMs) with a beta-binomial distribution were fitted using the *glmmTMB()* function from the *glmmTMB* package (Brooks et al. 2017). A beta-binomial distribution was chosen over a binomial distribution to account for overdispersion in the data. A separate GLM was fitted for each pollen fate, using the fate as the response variable (expressed in the model as a success/failure ratio; e.g., number of pollen grains deposited on stigmas/number of pollen grains not deposited on stigmas), and flower species (either *Solanum dulcamara* or *S. rostratum*) as the fixed effect. Estimated marginal means and confidence intervals for each species and pollen fate were obtained using the *emmeans()* function from the *emmeans* package (Lenth 2016).

Pollen deposition curves (Thomson & Plowright 1980) were analysed using generalized linear mixed-effects models (GLMMs) with a negative binomial distribution (log link, family = nbinom2), also fitted using *glmmTMB()*. A negative binomial distribution was chosen instead of a Poisson distribution to account for overdispersion, as pollen receipt data typically exhibit greater variance than expected under a Poisson model (Richards et al. 2009). The response variable was the number of pollen grains deposited on the stigma of a recipient flower. Fixed effects included recipient position (1-4, treated as numeric), flower species, the log-transformed duration (in seconds) of the floral visit, and the log-transformed number of pollen grains removed from the donor flower (calculated as B = F + G + H + D). Trial identity was included as a random intercept to account for repeated measures within trials.

To evaluate whether the shape of pollen deposition curves differed between species, two GLMMs were compared: one including species and recipient position as additive effects, and a second including their interaction. Model fit was assessed using the Akaike Information Criterion (AIC), with ΔAIC > 2 interpreted as evidence of better fit (Burnham & Anderson 2002). Model diagnostics, including checks for overdispersion and zero inflation, were performed using the DHARMa package (Hartig 2024).

Overall intercept and species-specific intercepts from pollen deposition curves were obtained using *emmeans()*. These represent the expected log-transformed number of pollen grains deposited on the first recipient flower, with other covariates held at their mean values. Additionally, estimated trends for continuous predictors (recipient position, visit duration, and number of donor pollen grains removed from the donor) were computed using *emtrends()*.

To assess how the number of sampled recipients influenced estimates of PTE, model predictions were used to estimate cumulative pollen deposition across a standardized sequence of 20 recipient flowers for each species, holding visit duration constant at its mean and fixing the number of released pollen grains to 100,000 for interpretability. Cumulative predicted values and their corresponding confidence intervals were summed across recipient positions for each species to estimate total pollen deposition. These values were then used to quantify how limiting the number of observed recipients (e.g., to four flowers) would underestimate total PTE.

All statistical analyses were conducted in R v.4.1.0 (R Core Team 2024) using *RStudio* (Posit Team 2024).

## Results

### Pollen fates and pollen transfer efficiency

The absolute numbers of pollen grains at each stage of the pollen transfer process are presented in Fig. 3. On average, *Solanum rostratum* flowers produced 28.97 % more pollen grains per flower than *S. dulcamara*. However, following a single visit, bees removed 37.77 % of the total pollen production in *S. dulcamara*, compared to just 13.55 % in *S. rostratum*—representing a nearly threefold higher proportional removal in *S. dulcamara*. Of the pollen removed, an average of 19.07 % in *S. dulcamara* and 62.99 % in *S. rostratum* was dislodged and fell to the ground. In addition, bees stored 65.92 % of the removed pollen in their corbiculae in *S. dulcamara*, and 27.65 % in *S. rostratum* (Fig. 4). These differences were significant for both pollen lost by dislodging (*Z* = 3.66, *P* = 0.001) and pollen stored in corbiculae (*Z* = –2.535, *P* = 0.011). On average, 14.01 % and 8.54 % of the removed pollen remained on the pollinator’s body at the end of a trial in *S. dulcamara* and *S. rostratum*, respectively, while only 0.86 % and 1.00 % were deposited on recipient stigmas (PTE). No significant differences were found between species in the proportion of pollen retained on the bee’s body (*Z* = –0.305, *P* = 0.760) or deposited on stigmas (*Z* = -1.223, *P* = 0.221). Estimated means and confidence intervals for each pollen fate by species, obtained from the GLMs, are provided in Appendix I.

**Figure 3.**
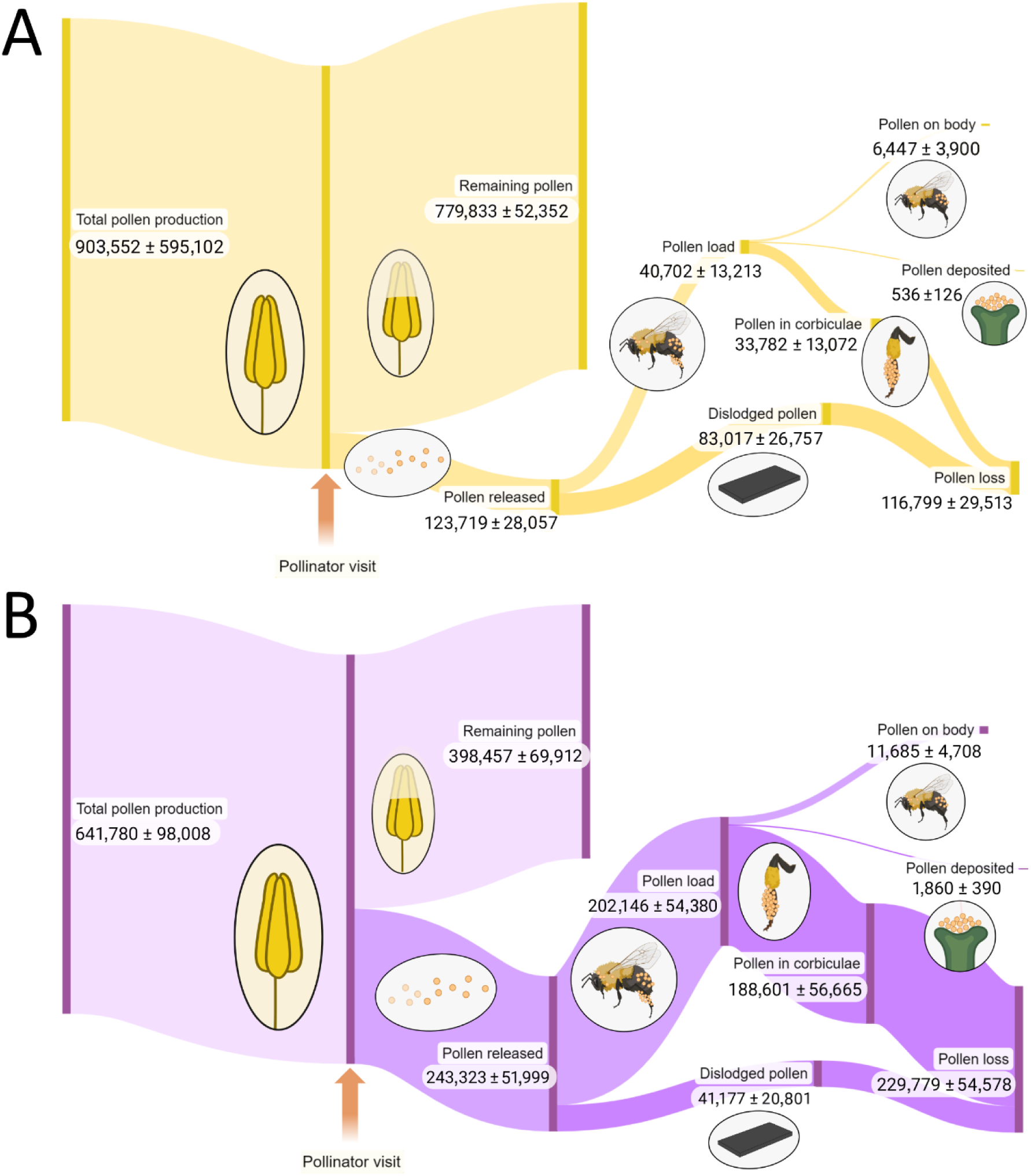
Diagram showing the fates of pollen of (A) *S. rostratum* and (B) *S. dulcamara*. The number of pollen grains for each stage of the pollen dispersal process is specified (mean ± S.E.). Created with BioRender.com

**Figure 4.**
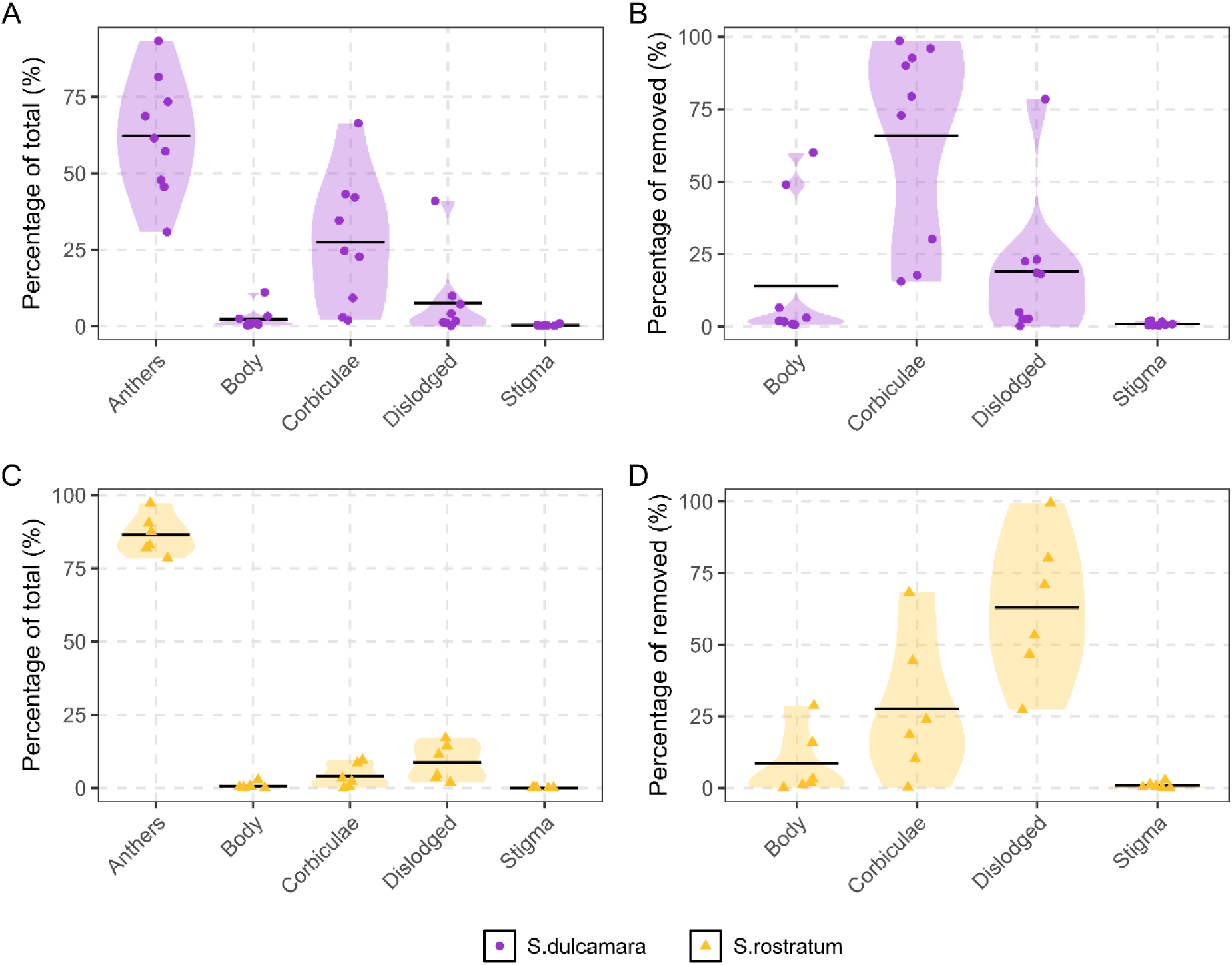
Pollen fates in *S. dulcamara* (A and B) and *S. rostratum* (C and D). On the left (A and C), the percentage of pollen grains in each measured variable is calculated from the total pollen production in a donor flower. On the right (B and D), the percentage of pollen grains reaching each location is calculated from the number of pollen grains removed by one bumblebee in a single visit to a donor flower. Black lines represent the mean.

### Pollen deposition curves

The additive model (excluding the interaction between recipient order and flower species) had a slightly lower AIC of 794.8, compared to 796.6 for the model including the interaction. As the AIC difference was less than 2 (ΔAIC = 1.8), both models were similarly supported; we therefore selected the additive model as the more parsimonious. Pollen deposition declined significantly with increasing recipient order (Fig. 5A; estimate = -0.339 ± 0.098, Z = -3.475, P < 0.001; Appendix II), indicating that most pollen was deposited on the first stigmas visited. Flowers of *S. rostratum* received significantly fewer pollen grains than those of *S. dulcamara* (Fig. 5A; estimate = -0.954 ± 0.310, Z = -3.082, P = 0.002; Appendix II). Longer visit durations (log-transformed) were associated with greater pollen deposition (Fig. 5B; estimate = 0.265 ± 0.110, Z = 2.399, P = 0.016; Appendix II). The number of pollen grains released from the donor flower (log-transformed) had no significant effect on pollen deposition (estimate = 0.155 ± 0.159, Z = 0.973, P = 0.330; Appendix II). To aid biological interpretation, Table 1 presents the overall and species-specific intercepts, as well as the estimated slopes for continuous predictors, representing the expected log-transformed number of pollen grains deposited on the first recipient flower, with covariates held at their mean values.

**Figure 5.**
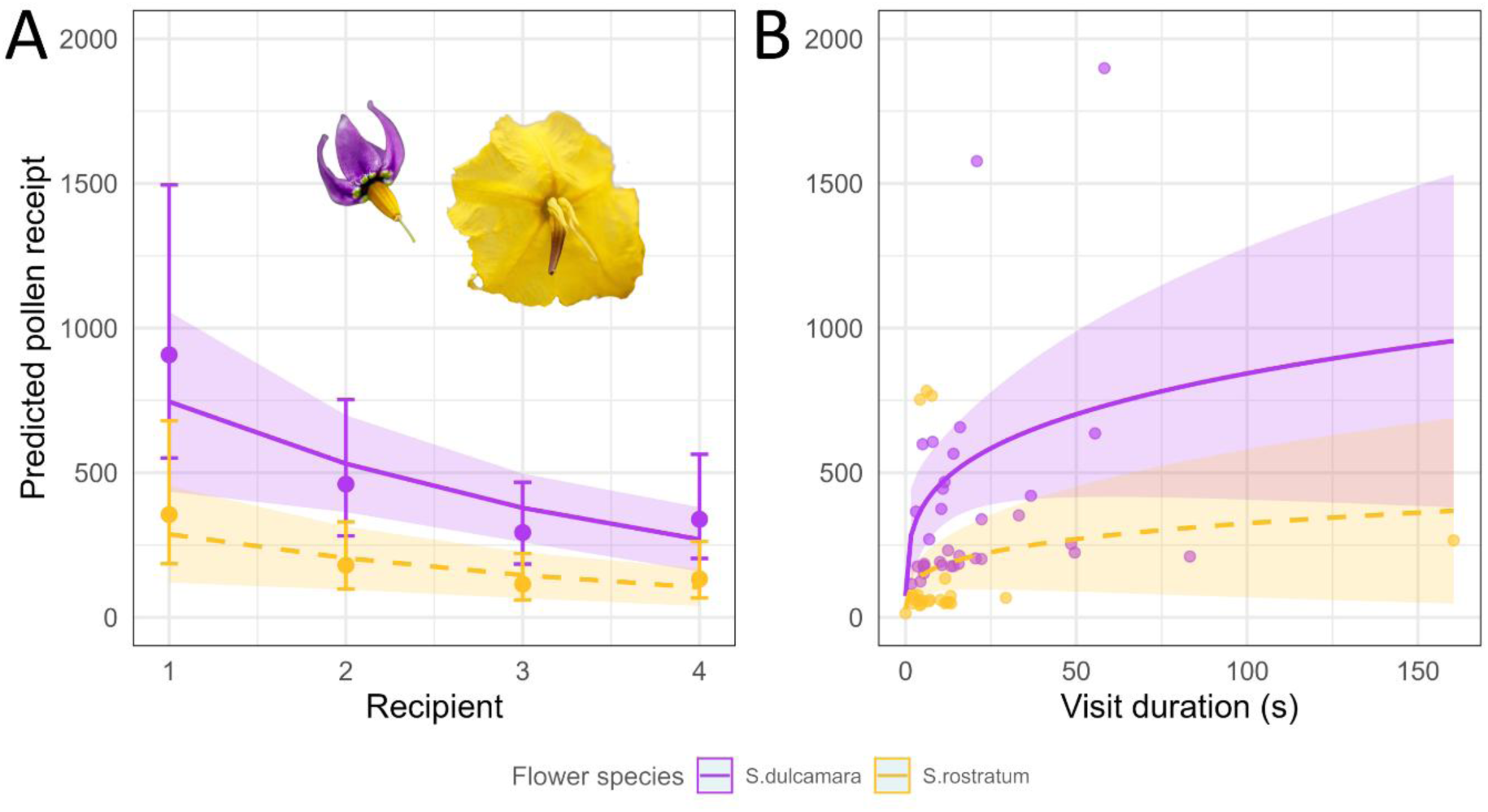
Pollen deposition on stigmas in relation to recipient position and visit duration for *S. dulcamara* and *S. rostratum*. (A) Predicted pollen receipt on stigmas across successive recipient positions (1–4) for each flower species. Solid and dashed lines represent model predictions and shaded regions represent 95% confidence intervals. Vertical bars and points represent estimated marginal means and standard errors for each recipient, obtained from a model treating recipient position as categorical variable. (B) Predicted pollen receipt as affected by visit duration in seconds for each flower species. Solid and dashed lines represent model predictions and shaded regions represent 95% confidence intervals. Points represent partial residuals (fitted values plus residuals on the link scale) for each species.

**Table 1.**
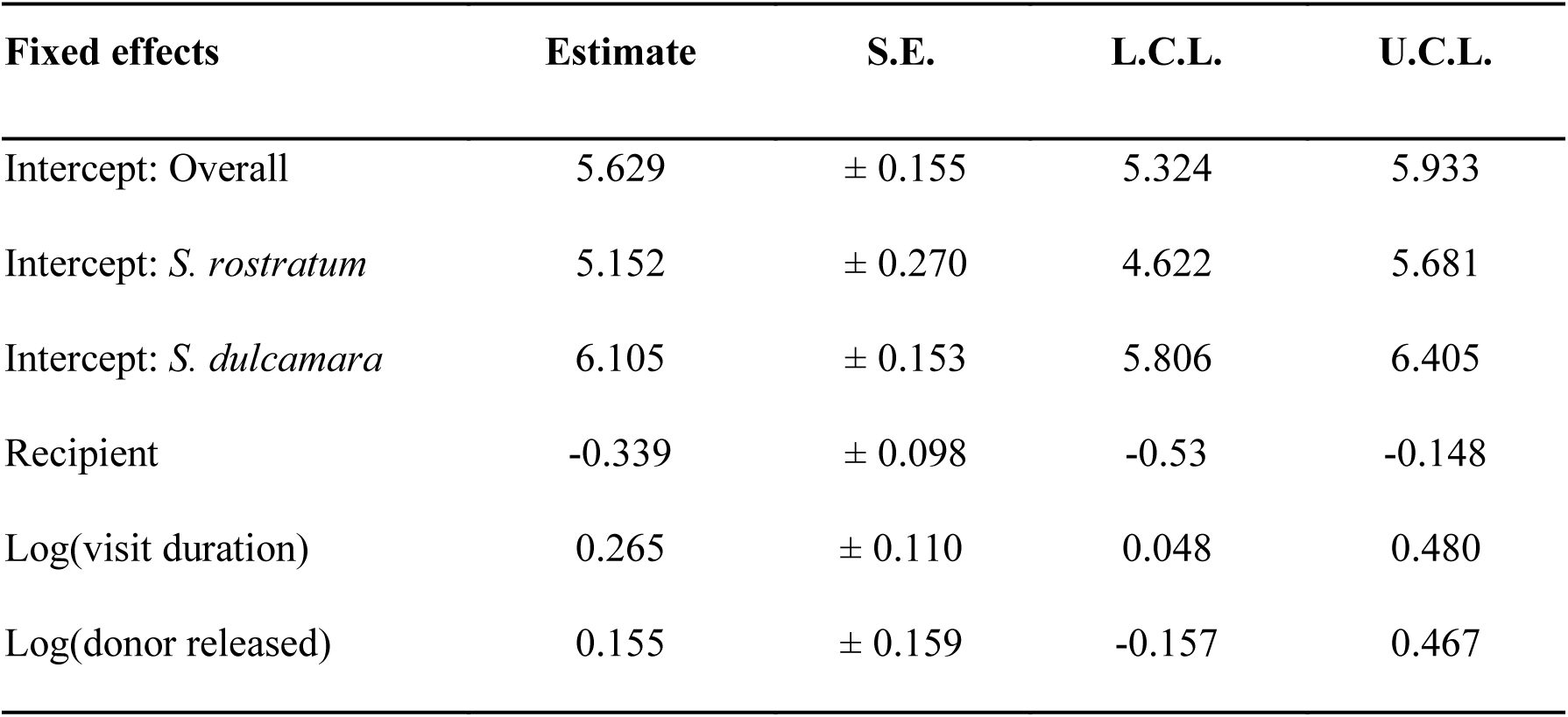
Estimated marginal means and trends from the selected negative binomial GLMM (log link) of pollen deposition on recipient stigmas. The overall intercept represents the expected log-transformed number of pollen grains deposited on the first recipient flower, with continuous covariates held at their mean values and species averaged. Species-specific intercepts represent expected values for *S. dulcamara* and *S. rostratum*, respectively. Estimated slopes (trends) are provided for recipient position, log-transformed visit duration (in seconds), and log-transformed number of pollen grains removed from donor flowers. S.E., standard error; L.C.L. and U.C.L., lower and upper 95% confidence limits, respectively.

### Underestimation of pollen transfer efficiency

In both *S. dulcamara* and *S. rostratum*, limiting sampling to four recipients underestimated pollen transfer efficiency (PTE) by ∼25.7% compared to estimates based on 20 recipients. PTE values increased with recipient number but at a decreasing rate, with underestimation falling below 2% by recipient 12 in both species. A summary and visualization of cumulative predicted pollen receipt and associated underestimation are provided in Appendices III and IV.

## Discussion

This study’s main findings can be summarised in three main points. First, both buzz-pollinated species of *Solanum* studied here had a similar or lower values of pollen transfer efficiency (PTE; Fig. 4) as those reported for other non-buzz pollinated plants that disperse pollen in individual pollen grain units (monads). Second, pollen deposition curves followed a classic exponential decay pattern, as in other animal-pollinated plants (Fig. 5). Finally, although both *Solanum* species studied showed a similar percentage of pollen loss, the pathway to this loss differed (Fig. 4). Pollen was either mostly lost on the ground (*S. rostratum*) or in the bees’ corbiculae (*S. dulcamara*), potentially reflecting the functional effects of different flower architectures. These findings and their implications are discussed below.

### Pollen transfer efficiency in buzz-pollinated flowers

Buzz pollinated species are thought to produce large numbers of pollen grains compared to nectar-producing relatives as a mechanism to offset the costs of active pollen collection by floral visitors (De Luca & Vallejo-Marín 2013; Vallejo-Marín & Russell 2024). The average pollen production per flower is indeed high in *S. dulcamara* (> 600,000) and *S. rostratum* (> 900,000) (Fig. 3), and comparable to other buzz-pollinated plant species (Vallejo-Marín et al. 2009; Vallejo-Marín et al. 2014; Vallejo-Marín et al. 2022). In this experiment, visitation by a generalist, buzz-pollinating bumblebee resulted in similar pollen transfer efficiency (PTE) between both *Solanum* species, with mean values of 0.86 % and 1.00 % for *S. rostratum* and *S. dulcamara*, respectively (Fig. 4). The use of only four recipients likely underestimated PTE in our experiment. We estimated that, were we to measure pollen receipt in 20 flowers instead of four, PTE values would have increased by about 25 %. Pollen transfer efficiency has only been rarely quantified for buzz pollinated plants, but low PTE characterizes two previously studied species. *Rhexia virginica* (Melastomataceae), an herbaceous species distributed in eastern North America, has a PTE of 0.92 % (Supplementary material from Johnson & Harder 2023), while *Senna reticulata* a neotropical tree, has a much lower PTE (0.05 %) (Supplementary material from Johnson & Harder 2023). These values of PTE in buzz pollinated species might suggest that active pollen collection by bees indeed impacts the realized efficiency of pollen transfer as they are highly effective in grooming and have specialized structures for collecting and retaining large amounts of pollen (Thorp 2000). However, further work comparing PTE in buzz pollinated taxa and their closest non-buzz pollinated relatives in similar field settings is required to establish if low PTE indeed characterizes buzz-pollinated flowers. Furthermore, PTE can vary among different species of floral visitors, and this is an area that also requires additional study.

### Pollen removal, flower morphologies and mating systems

Percentage of pollen removed by a single pollinator visit in other buzz pollinated species ranges from 10-70 % (*Dodecatheon conjugens:* 20-70 %, depending on flower age; Harder & Barclay 1994; *Rhexia virginica:* 10-43 %; Larson & Barrett 1999; *Senna reticulata:* 25-75 %; Snow & Roubik 1987; *Erica tetralix* and *Vaccinum myrtillus*: 28-60 %; Moquet et al. 2017), and is similar or lower than in non-buzz-pollinated species such as *Aconitum septentrionale* (44-61 %; Thøstesen & Olesen 1996), *Aralia hispida, Mertensia paniculata, Aconitum delphinifolium, Pedicularis bracteosa, Pedicularis contorta and Lupinus sericeus* (79.6 %, 62.2 %, 37.7-61.4 %, 48.7 %, 43.5 %, 18.9 %, respectively; Harder 1990) *Malus domestica* and *Prunus dulcis* (40-50 %; Thomson & Goodell 2001), and *Calluna vulgaris* (≤ 88 %; Moquet et al. 2017). The small-flowered *S. dulcamara* produces fewer pollen grains than *S. rostratum* but a higher number (and percentage) of pollen grains are removed during a single visit by a bumblebee (37.77 % pollen removed per visit against 13.55 % for *S. rostratum*). Fewer visits by a pollinator might therefore be needed to empty a *S. dulcamara* flower. However, pollen inside the anther may not all be available to be removed at once, and it appears that some buzz pollinated flowers present a staggered release of pollen as the flower ages (Harder & Barclay 1994; Kemp & Vallejo-Marín 2021). The extent to which pollen becomes available simultaneously or in a staggered fashion in buzz pollinated species is not well understood. Regardless of the mechanism, the high rates of pollen removal observed in *S. dulcamara* are characteristic of other small-flowered, non-heterantherous taxa in *Solanum* Section Androceras (e.g., *S. fructu-tecto*), and this has been interpreted as a consequence of a shift in the mating system towards increased selfing (Kemp & Vallejo-Marín 2021) or perhaps lower visitation rates. The mating system of *S. dulcamara* is presently unknown, but this self-compatible species sometimes sets fruit autonomously (M. Vallejo-Marín, Uppsala University, Sweden, pers. obs). In contrast, although *S. rostratum* is also self-compatible, it has a high outcrossing rate (0.70-0.90) (Vallejo-Marín et al. 2013; Morra-Carrera et al. 2019) and rarely sets fruits autonomously, at least in the native range. In addition, *S. dulcamara* presents anthers joined in a cone, while in *S. rostratum,* the anthers are relatively free to move from one another (Fig. 2B). Experimental work has demonstrated that joined anther cones facilitate the transmission of vibrations during buzzing and increase pollen release (Vallejo-Marín et al. 2022). Thus, the difference in the proportion of pollen removed per visit could be associated to different floral morphologies, visitation rates, and mating systems in *S. dulcamara* and *S. rostratum.* For outcrossing species visited regularly by pollinators, a more restricted pollen schedule might allow them to dispense pollen more efficiently or to more flowers (Harder & Thomson 1989; Harder & Wilson 1994; Kemp & Vallejo-Marín 2021).

### Patterns of pollen loss

It is likely that the estimates of pollen loss presented in Fig. 4 do not capture the full extent of the actual losses. Transport loss (pollen lost on the ground or in the air after a bee left the donor flower) was not measured in the trials, leading to an underestimation of dislodged pollen (D). Flight distance between the four flowers presented were also smaller than in field conditions, which could reduce the capacity for grooming in the air, and therefore, the estimations of pollen lost in the corbiculae (F). However, in-flight grooming appears to occur mostly while the bee hovers near the flower (MVM pers. obs.), so perhaps travel distance has less of an effect on grooming capacity. Such considerations highlight the challenges of experimentally studying pollen fates and suggest caution when comparing our results to field studies in other species. However, an interesting pattern appeared when comparing both species studied here. Although they exported a similar proportion of pollen grains to stigmas, the pathway to pollen loss differed: in *S. dulcamara*, pollen was mostly lost to bees collecting it and placing it in their corbiculae, while in *S. rostratum,* pollen was primarily lost by falling on to the ground (Fig. 4), meaning that this dislodged pollen did not contribute to the fitness of either bee or plant. What is the reason for these contrasting pollen fates in these two species? With only two species studied, it is difficult to generalize. A potential explanation lies in that the bees can curl their body in proximity of the anther pores of *S. dulcamara* while buzzing the flower. The fused anther cone of *S. dulcamara* then works as a single pollen dispensing unit that places pollen grains in the ventral region of the bee where it can be groomed and placed in the corbiculae (Nevard & Vallejo-Marín 2022). In contrast, in heterantherous *S. rostratum*, the bee cannot work simultaneously both feeding and pollinating anthers, and it usually focuses its attention on the feeding anthers. The feeding anthers contact the bee on the ventral side, while the pollinating anther contacts the bee on the dorsal side of the abdomen, which is harder for bees to groom. The body of the bee often contacts the terminal pores of the feeding anthers, but the distance between the bee’s body and the terminal pore of the pollinating anther depends both on the positioning of the bee and on the relative size of bee and flower (Solís-Montero & Vallejo-Marín, 2017). Because approximately 50 % of the total pollen production in these flowers resides on the single pollinating anther (Vallejo-Marín et al. 2009; Vallejo-Marín et al. 2014), flower handling and bee-flower size matching can then affect significantly whether the pollen reaches the bee’s body or falls to the ground. Although this was not experimentally quantified, it is expected that much of the pollen loss occurring during buzz pollination in *S. rostratum* was due to mismatches between the size of the bee and the flower (Solís-Montero & Vallejo-Marín, 2017). All else being equal, a bee of similar characteristics to the ones studied here might benefit more from a single visit to a species like *S. dulcamara* than one like *S. rostratum.* Of course, differences in total pollen production, number of flowers per plant, increased visitation time and repeated visits to a flower might help offset the increased difficulty of gathering pollen from flowers with a more complex morphology as in *S. rostratum*.

### Pollen deposition curves

The results presented here demonstrated that pollen deposition curves (i.e., the pattern of pollen deposition on the stigmas of consecutive recipients) for the buzz-pollinated *Solanum* species studied followed a typical exponential decay pattern (Fig. 5), at least within the range of evaluated recipients, as reported for other species (Thomson & Plowright 1980; Mitchell et al. 2013; Santa-Martinez et al. 2021). Determining if this curve exhibits a leptokurtic pattern would imply evaluating a longer recipient sequence. Specifically, the amount of pollen deposited on stigmas decreased by 33 % with each successive recipient for both *S. dulcamara* and *S. rostratum*. These percentages are on the higher end of the spectrum for pollen carryover curves, which ranges from 50 % to 0.5 % (see Tab. 1 in Robertson 1992), indicating a relatively steep decline in deposited pollen with each successive recipient. Generally, grooming pollinators such as bees cause steeper deposition curves, indicating less extensive pollen carryover, compared to pollinators like hummingbirds and hawkmoths (Robertson 1992; Castellanos et al. 2003; Mitchell et al. 2013). However, deposition curves have not been studied in many other buzz-pollinated plants, highlighting a gap in our understanding of these systems.

### Implications of steep pollen deposition curves for gene flow

Pollen deposition curves are influenced by both the amount of pollen that lands on the pollinator’s body and the intensity of its grooming before visiting the next flower. Buzz-pollinated *Solanum* flowers, which produce and gradually release large amounts of pollen as a reward for pollinators, tend to encourage more intense grooming compared to nectar-producing, non-buzz-pollinated flowers (Harder 1990; Johnson & Harder 2023). This intense grooming results in steeper pollen deposition curves (Holmquist et al. 2012), potentially influencing gene flow patterns (Santa-Martinez et al. 2021). Consequently, if buzz-pollinated flowers generally exhibit steeper pollen deposition curves, they might experience more restricted gene flow compared to non-buzz-pollinated species offering other rewards, such as nectar. While insect-pollinated species often display higher levels of population genetic structure (Dellinger et al. 2022; Gamba & Muchhala 2020; Gamba & Muchhala 2023), the specific impact of buzz pollination on genetic structure remains underexplored.

## Conclusion

Empirical studies of pollen fates, though complex and time-consuming, provide relevant insights into plant reproductive success. While controlled laboratory experiments may not fully reflect the complexity of natural systems, they offer a foundational understanding of pollen dynamics and patterns in plant reproduction. Additionally, the data they provide are often unattainable under field conditions. This study found significant differences in pollen fates, but not in the steepness of deposition curves, between *S. dulcamara* and *S. rostratum*, highlighting the potential impact of floral architecture on pollen fates, even among species visited by the same buzz-pollinating bee. Although future studies including longer visitation sequences are needed to more precisely estimate PTE, our findings suggest that Pollen Transfer Efficiency (PTE) in pollen-rewarding, buzz-pollinated species is comparable or lower than the average of 2.3% reported for nectar-rewarding species with single pollen units (Johnson & Harder 2023). Further studies on PTE of a wider range of species with diverse morphological traits and reward types will help determining how flower morphology can shape pollen fates.

## Acknowledgments

The authors thank D. Scaccabarozzi and L. van Kolfschoten for their technical support and advice throughout the development of this work. The authors are also grateful to the Research Experience for Peruvian Undergraduates (REPU) Program and the University of Lausanne exchange program for enabling A.V.C., E.M., and B.G.S. to carry out this project at Uppsala University. A.V.C., E.M., B.G.S., and M.V.M conceived the study. A.V.C., E.M. and B.G.S. conducted the experiments, collected the data and performed the analyses. A.V.C., E.M., B.G.S., and M.V.M wrote and revised the manuscript. M.V.M supervised the study and acquired funding. This work was supported by start-up funds from the Plant Ecology and Evolution program at Uppsala University, awarded to M.V.M.

## Data availability

Pollen transfer data used for the analyses and the R script will be made available upon acceptance.

## Appendices

**Appendix I.** Estimated marginal means of the proportion of pollen in each fate by species, obtained from beta-binomial GLMs. The table also summarizes the effects of flower species (S. dulcamara, S. rostratum) on the proportion of pollen allocated to four fates. S.E., standard error; L.C.L. and U.C.L., lower and upper 95% confidence limits, respectively; Z, z-value; P, p-value. N = 15, residual degrees of freedom = 12.

| Pollen fate | Species | Estimated mean | L.C.L. | U.C.L. | Z | P |
| --- | --- | --- | --- | --- | --- | --- |
| Body | *S.dulcamara* | 0.133 | 0.064 | 0.255 | -0.305 | 0.760 |
|  | *S.rostratum* | 0.115 | 0.048 | 0.252 |  |  |
| Corbiculae | *S.dulcamara* | 0.661 | 0.478 | 0.805 | -2.535 | 0.011 |
|  | *S.rostratum* | 0.281 | 0.133 | 0.497 |  |  |
| Dislodged | *S.dulcamara* | 0.212 | 0.111 | 0.368 | 3.266 | 0.001 |
|  | *S.rostratum* | 0.668 | 0.460 | 0.826 |  |  |
| Stigma | *S.dulcamara* | 0.011 | 0.007 | 0.017 | -1.223 | 0.221 |
|  | *S.rostratum* | 0.007 | 0.004 | 0.013 |  |  |

**Appendix II.** Summary of the statistical analysis evaluating the effects of recipient order, flower species (*S. dulcamara*, *S. rostratum*), log-transformed visit duration, and log-transformed number of donor pollen grains removed on pollen deposition. Model estimates were obtained from a GLMM with trial identity as a random effect, and recipient order, flower species, and visit duration as fixed effects. S.E., standard error; Z, z-value; P, p-value. N = 60, residual degrees of freedom = 53.

| **Fixed effects** | **Estimate** | **S.E.** | **Z** | **P** |
| --- | --- | --- | --- | --- |
| Intercept | 4.310 | ± 2.010 | 2.144 | 0.032 |
| Species: *S. rostratum* | -0.954 | ± 0.310 | -3.082 | 0.002 |
| Recipient | -0.339 | ± 0.098 | -3.475 | 0.001 |
| log(visit duration) | 0.265 | ± 0.110 | 2.399 | 0.016 |
| log(donor released) | 0.155 | ± 0.159 | 0.973 | 0.330 |

**Appendix III.**
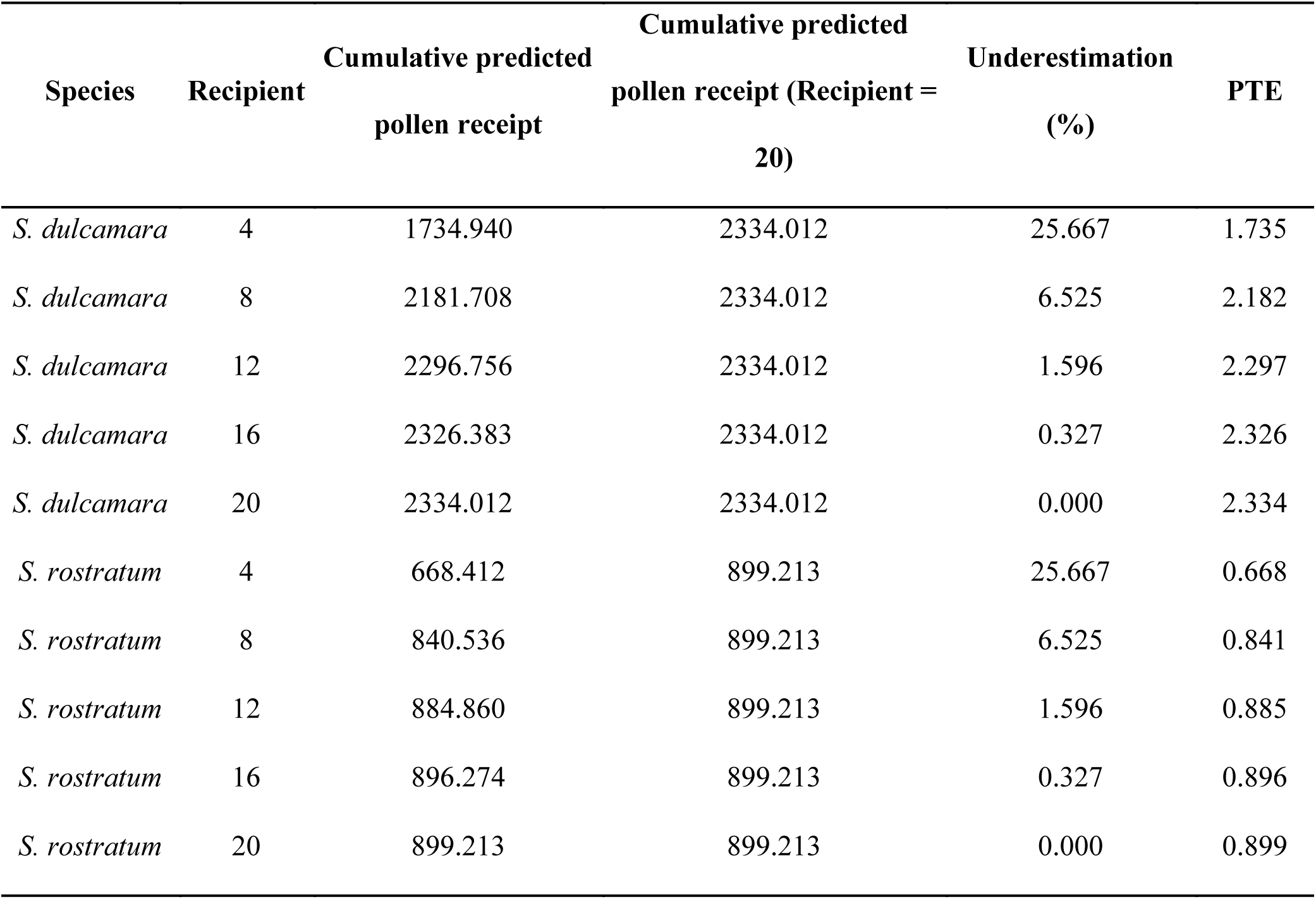
Summary of cumulative predicted pollen receipt and corresponding underestimation across increasing numbers of sampled recipients for *S. dulcamara* and S. rostratum. Cumulative deposition values are expressed per 100,000 pollen grains removed. Percent underestimation indicates the proportion of total predicted pollen not captured when sampling is limited to a given number of recipients.

**Appendix IV.**
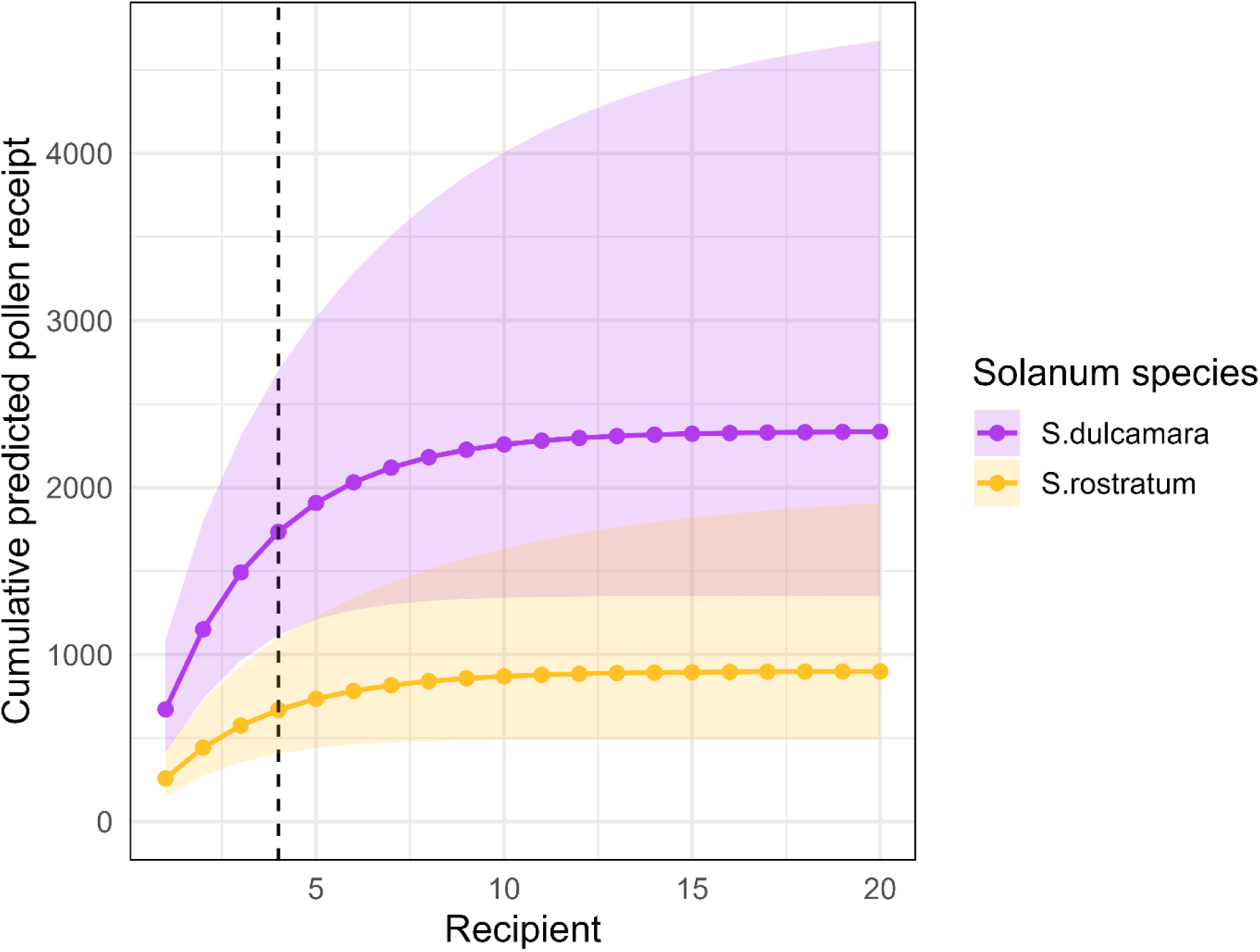
Cumulative predicted pollen deposition on recipient stigmas across consecutive flowers in a bee’s visit sequence for S. dulcamara and S. rostratum. Lines show model-predicted cumulative deposition based on a negative binomial GLMM, assuming a constant pollen release of 100,000 grains and average visit duration. Shaded areas represent 95% confidence intervals. The dashed vertical line indicates the fourth recipient, which corresponds to the maximum number of recipients sampled in the experiment.

